# A snapshot of brain and cognition in healthy mid-life and older adults

**DOI:** 10.1101/2022.01.20.476706

**Authors:** Léonie Borne, Michelle K. Lupton, Christine Guo, Philip Mosley, Robert Adam, Amelia Ceslis, Pierrick Bourgeat, Amir Fazlollahi, Paul Maruff, Christopher C. Rowe, Colin L. Masters, Jurgen Fripp, Gail A. Robinson, Michael Breakspear, for the Prospective Imaging Study of Ageing, the Alzheimer’s Disease Neuroimaging Initiative, the Australian Imaging Biomarkers and Lifestyle flagship study

**Author notes:** Data used in preparation of this article were obtained from the Prospective Imaging Study of Ageing (PISA): Genes, Brain and Behaviour database (qimrberghofer.edu.au/study/prospective-imaging-study-of-ageing), funded by the National Health and Medical Research Council (NHMRC) of the Australian Government. The PISA investigators contributed to the design and implementation of PISA and/or provided data but did not participate in analysis or writing of this report. Data used in preparation of this article were obtained from the Alzheimer’s Disease Neuroimaging Initiative (ADNI) database (adni.loni.usc.edu). As such, the investigators within the ADNI contributed to the design and implementation of ADNI and/or provided data but did not participate in analysis or writing of this report. A complete listing of ADNI investigators can be found at:http://adni.loni.usc.edu/wp-content/uploads/how_to_apply/ADNI_Acknowledgement_List.pdf. Data used in the preparation of this article was obtained from the Australian Imaging Biomarkers and Lifestyle flagship study (AIBL) made available at the ADNI database (www.loni.usc.edu/ADNI). The AIBL researchers contributed data but did not participate in analysis or writing of this report. AIBL researchers are listed at www.aibl.csiro.au.

## Abstract

Deficits in memory are seen as a canonical sign of ageing and a prodrome to dementia in older adults. However, the nature of cognitive and brain changes across a wider aperture of adulthood is not well known. We quantify the relationship between cognitive function and brain morphology from mid-life to older adulthood, and the influence of age, sex, amyloid and genetic risk for dementia. We analyzed three observational cohorts (PISA, AIBL, ADNI) with cognitive, genetic and neuroimaging measures comprising a total of 1570 healthy mid-life and older adults (mean age 72, range 49-90 years, 1330 males) and 1365 age- and sex-matched adults with mild cognitive impairment or Alzheimer’s disease. Among healthy adults, we find robust modes of co-variation between regional sulcal width and multidomain cognitive function that change from mid-life to the older age range. The most prominent cortical changes in mid-life are predominantly associated with changes in executive functions, whereas they are most strongly associated with poorer memory function in older age. These cognitive changes are accompanied by an age-dependent pattern of sulcal widening. Amyloid exerts a weak, but significant, influence on cognition, but not on sulcal width. The *APOE* ɛ4 allele also exerts a weak influence on cognition, but only significantly in the (larger and older) AIBL cohort. These findings provide new insights into brain and cognition in mid-life and older adults, suggesting that cognitive screening in mid-life cohorts should encompass executive functions as well as memory.

## Introduction

Ageing is accompanied by substantial cognitive and neuronal changes that reflect the direct influence of neuronal loss and accumulating risk factors, such as chronic cardiovascular and metabolic disease. These age-related changes in brain and cognition are challenging to identify for several reasons. First, their relatively slow development makes them difficult to detect in a longitudinal study that only spans a few years. Second, the cognitive and anatomical variability between individuals may be larger than the smaller age-related differences across a cross-sectional study (Raz et al., 2005). Healthy aging is also difficult to discern from the influence of neurodegenerative diseases such as Alzheimer’s disease (AD) (Doan et al., 2017; Lorenzi et al., 2015; Raz et al., 2010). Indeed, the preclinical stage of AD may begin several decades before its diagnosis (Morris, 2005) hence commencing in the mid-life decades. The accurate identification of this early stage would open new diagnostic and therapeutic windows (Barnett et al., 2014; Dubois et al., 2016) but requires delineation from healthy ageing across a broad sweep of adulthood.

Healthy aging in older adults is accompanied by both grey and white matter changes (Gunning-Dixon et al., 2009; Hugenschmidt et al., 2008; Salat et al., 2005). Studies showing sex differences in age-related white matter changes have had inconsistent findings (Cox et al., 2016; Hsu et al., 2008). Some regions, such as the prefrontal cortex, appear particularly sensitive to these changes, while others, such as the medial temporal cortex or the hippocampus, are relatively preserved (Good et al., 2001; Salat et al., 2004). These changes manifest specific cognitive changes. For example, lateral prefrontal changes are associated with a decline in executive function (Burzynska et al., 2012; Westlye et al., 2011).

Neurodegenerative changes in the early stages of Alzheimer’s disease occur predominantly in the hippocampus, precuneus and medial temporal lobe, often manifesting as amnesia (Coupé et al., 2019; He et al., 2007; Jack et al., 1992; Mu and Gage, 2011; Scheltens et al., 1992). Several risk factors for AD have been identified, including genetic factors such as the *APOE* ɛ4 allele (Saunders et al., 1993) and environmental factors such as fewer years of education (Livingston et al., 2020). AD progression is linked to molecular mechanisms, associated with the formation of amyloid plaques (Chiti and Dobson, 2006) early in the disease (Sperling et al., 2011). While these disparate features of AD are well documented, their emergence from the healthy mid-life brain is not understood.

There is a gap in the literature regarding cognitive and brain changes in mid-life (Raz et al., 2010). Large multimodal studies of AD, such as the Alzheimer’s Disease Neuroimaging Initiative (ADNI) (Mueller et al., 2005) and the Australian Imaging, Biomarkers and Lifestyle (AIBL) (Ellis et al., 2009) focus on the decades for peak onset of AD, i.e. 60-80 years, after the first neurobiological changes of AD, which likely begin several decades earlier (Dubois et al., 2016; Morris, 2005). The Prospective Imaging Study of Ageing: Genes, Brain and Behaviour (PISA) (Lupton et al., 2021), is a multimodal study including amyloid PET scans that studies midlife ageing (mean age 61, range 49-73), with a focus on individuals at high genetic risk of developing AD (*APOE* ɛ4 positive as well as those with high polygenic risk scores, AD-PRS). An integrative analysis of these large cohort studies would thus enable a snapshot of brain and cognition across a wide aperture of mid-life and older adulthood.

Here, we investigate and compare the relationship between cognition and brain morphology in cross-sectional data for two age ranges: one (using PISA) to focus on the mid-life age range for the preclinical stage of AD and the other (using ADNI and AIBL) focussed on an older age-range corresponding to the typical onset of AD. We use a multivariate method, the canonical Partial Least Square (Wegelin, 2000) which enables a global and unbiased analysis of the relationship between brain and behaviour in these three cohorts. For the measure of brain anatomy we use sulcal width (SW) derived from structural magnetic resonance imaging (sMRI). SW is a promising marker for the sensitive detection of disease which has been shown to be more accurate than cortical thickness (CT) in differentiating mild cognitive impairment (MCI) and AD from healthy people (Bertoux et al., 2019). We then examine how brain-cognition relationships are modified by age, sex, cortical amyloid and genetic risk factors for AD across these three cohorts. To explore putative relationships between ageing and neurodegeneration, we benchmark the analyses of all three healthy cohorts against clinical cohorts of age- and sex-matched persons with MCI or AD.

## Results

We analyzed cognitive, neuroimaging (MRI, PET) and genetic data from 1570 healthy adults (healthy cohorts, HC) and 1365 adults with MCI or AD (clinical cohorts, CC), drawn from 3 multimodal databases; PISA, ADNI and AIBL. While the age ranges of these cohorts mutually overlap, the PISA cohort spans a younger mid-life cohort than AIBL and ADNI (Fig. 1). To integrate brain and cognitive data, we used partial least squares (PLS), a multivariate method that identifies modes of covariation between two multivariate data set, here regional SW and multidomain neurocognitive scores (see Methods). Nonparametric testing was performed to identify robust modes of covariation (at *p*<0.05).

**Figure 1:**
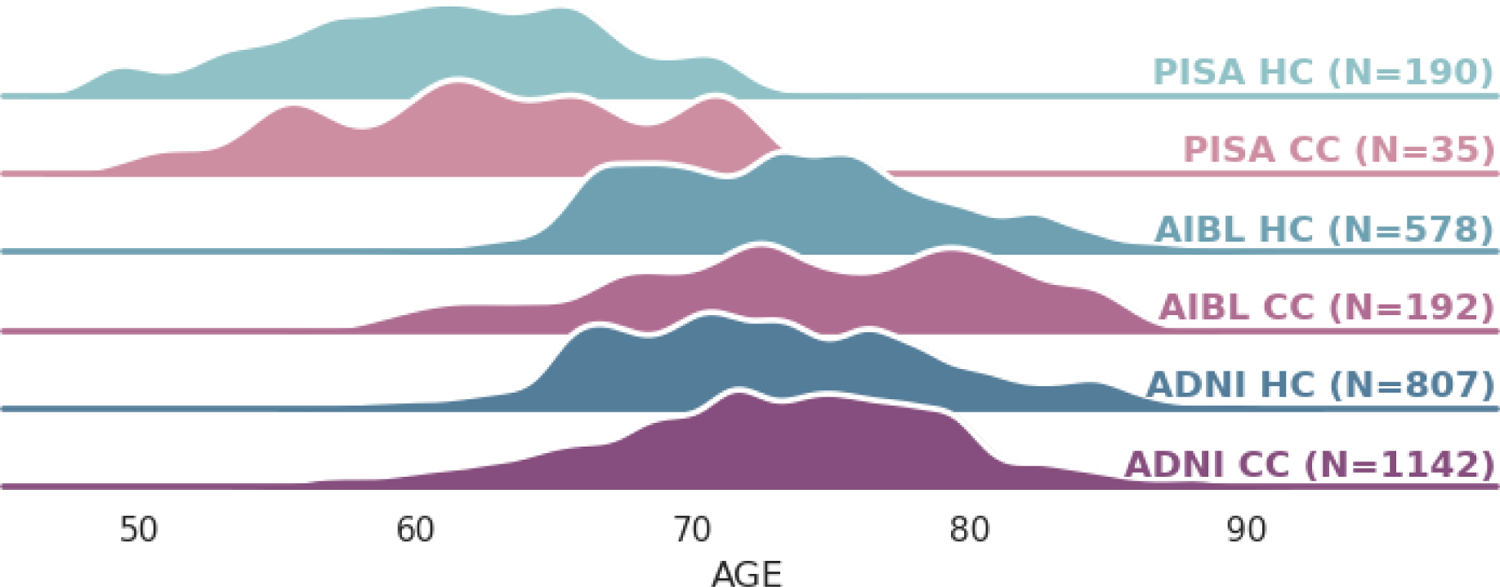
**Age distribution for the three databases (PISA, AIBL and ADNI)**, each comprising a healthy cohort (HC) and a clinical cohort (CC) containing participants with MCI or AD. HC and CC participants in each database were matched for age and sex.

### 1. Brain-behaviour modes in mid-life PISA participants

Application of PLS to the 190 healthy PISA HC participants, using SW as the anatomical measure, yields a single robust mode (1st mode, *p*<0.001, *z*-covariance=5.49; 2nd mode, *p*>0.99). Using cortical thickness (CT) as the brain measure also yields a single mode, but with less covariance explained (1st mode, *p*=0.0060, *z*=3.07; 2nd mode, *p*>0.99). We hereafter focus on the SW-derived mode as it explains greater than 50% more brain-behaviour covariance in all PLS analyses (see Supp. Fig. 2 for the CT-derived PLS analyses).

**Figure 2:**
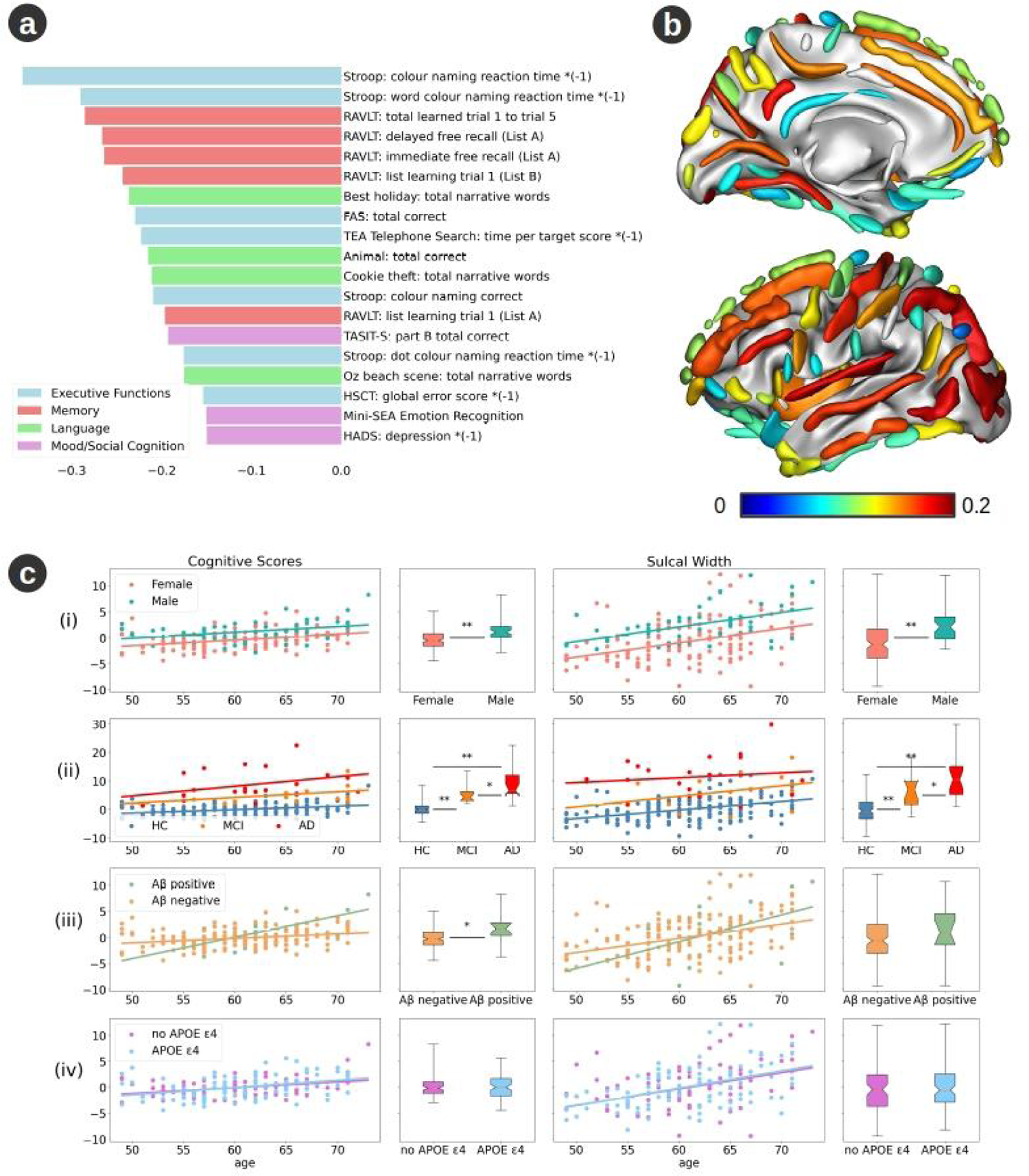
Robust PLS mode trained on PISA healthy cohort. (a) Mean loading of all reliable cognitive scores, colour-coded by domain. Larger (negative) projections denote poorer performance. (b) Mean loading of all reliable cortical sulci. Higher projections denote wider sulci. (c) Effect of age on PISA brain-behaviour mode. Cognitive (left) and sulcal width (right) projections of the single robust mode. More positive projections denote poorer cognitive performance and wider sulci, respectively. (i) Healthy male (cyan) and female (orange) participants, (ii) Healthy amyloid positive (orange) or negative (green) participants, (iii) Healthy cohort (HC, blue) and participants with MCI (orange) or AD (red), (iv) Healthy participants with (blue) or without (purple) the allele 4 of the *APOE* gene. Lines and scatter plots show the evolution of the projections as a function of age. The box plots show the distribution of the cognitive and brain projections by group. The box extends from the lower to upper quartile values of the data, with a notch at the median. The whiskers extend from the box from the minimum to the maximum values. The influence of groups on the projection is evaluated with an ANCOVA controlling for age and sex. Significant at *p* < 0.05*; *p*< 0.001**.

This robust PLS mode loads across all three main cognitive domains, but with the strongest influence of the executive function tests (Fig. 2a). The brain projection of the mode (SW) loads most strongly onto the occipital lobe, the intraparietal sulcus, the posterior inferior temporal sulcus, the posterior lateral fissure and the sub-parietal sulcus (Fig. 2b, Supp. Fig. 1). Both the cognitive and brain projections are strongly correlated with the age of the participants (SW, *p*=6.1e-9; cognition, *p*=8.7e-7). The projections also differ as a function of sex, when controlling for age (SW, *p*=1.2e-6; cognition, *p*=1.6 e-6; Fig. 2c).

We applied this PLS model - trained on healthy participants - to the 35 age- and sex-matched MCI and AD PISA participants (Fig. 2c). The brain and cognitive projections are greater with disease stage – i.e. are significantly higher for AD than for MCI and in turn for MCI compared to HC. Furthermore, the age-dependent slope differs between healthy and AD subjects for the cognitive projection (*p*=0.022), with a faster age-related decline across the AD participants. Among healthy participants, the cognitive projection is significantly influenced by the presence of amyloid (*p*=0.0019) but not by the *APOE* ɛ4 allele (*p*=0.69). Specifically, amyloid positive participants have significantly more severe age-related cognitive changes, with a significantly different age-dependent slope compared to amyloid negative participants (*p*=6.6e-04). The SW projection is not influenced by either of these factors (amyloid, *p*=0.85; *APOE*, *p*=0.55; Fig. 2c).

### 2. Brain-behaviour modes in older adult AIBL participants

Application of PLS to the 573 healthy older adult AIBL participants also yields a single robust mode (*p*<0.001, *z*=6.60). As with the PISA data, this single SW-derived mode loads across all three main cognitive domains but, notably, with strongest affinity for the memory domain, not executive function (Fig.3a). The brain projection loads most strongly over the superior temporal sulcus, the posterior lateral fissure, the inferior frontal sulcus, the occipital lobe and the intermediate frontal sulcus (Fig. 3b; Supp. Fig. 1). Interestingly, the first mode is not robust when using CT as the brain measure (*p*=0.11).

**Figure 3:**
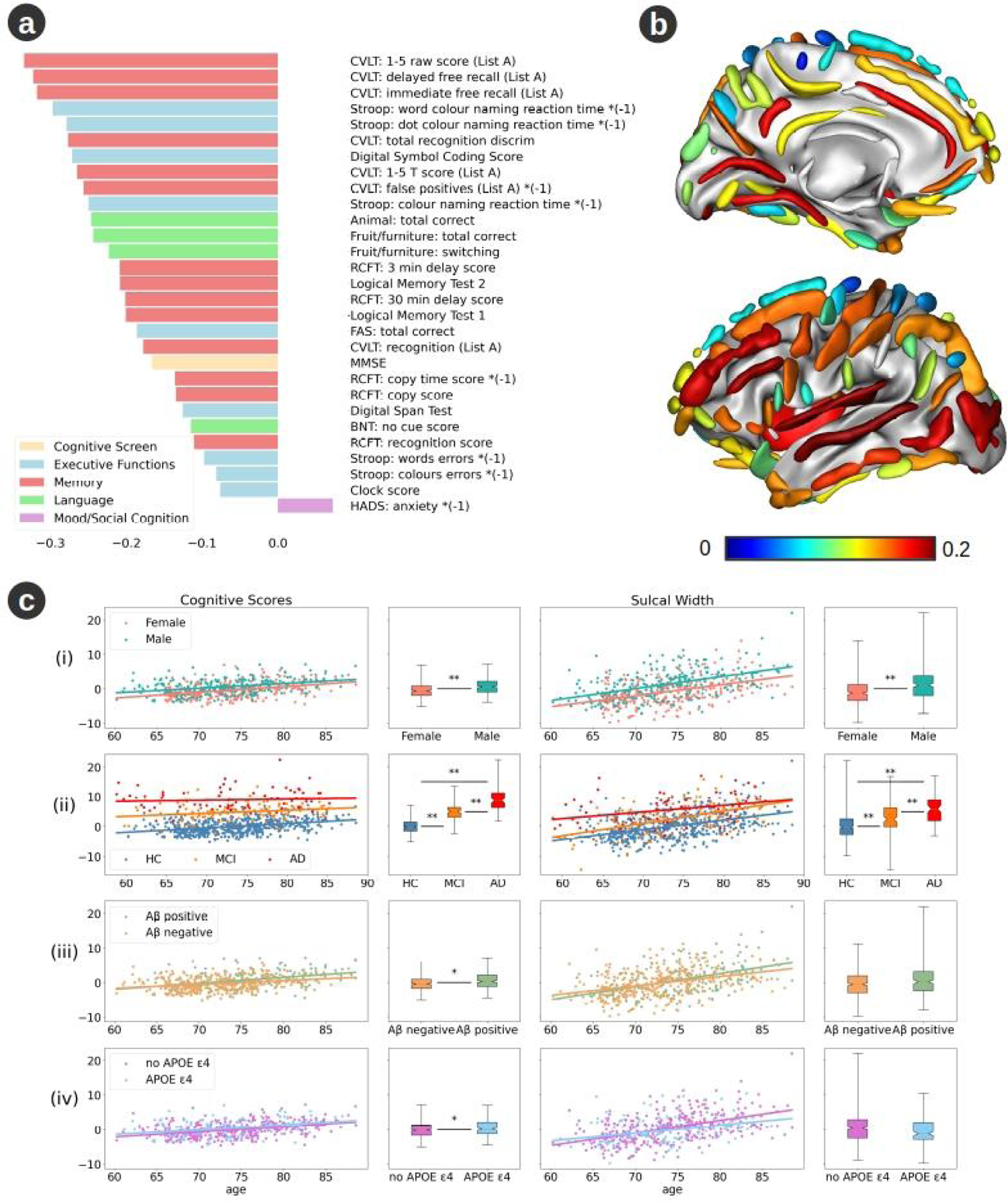
Robust PLS mode trained on AIBL healthy cohort. (a) Mean loading of all reliable cognitive scores. (b) Mean loading of all reliable cortical sulci. (c) Effect of age on AIBL brain-behaviour mode. Panels and legend as per Figure 2.

As with the PISA data, projections of both cognition and SW correlate significantly with age (SW, *p*=9.7e-29; cognition, *p*=1.0e-19) and sex, when controlling for age (SW, *p*=4.2e-13; cognition, *p*=9.0e-11, Fig. 3c). The PLS model trained on healthy participants, controlling for age and gender, again loads more strongly onto AD than MCI participants, and more strongly for MCI than for HC participants (Fig 3c). Furthermore, the age-dependent slope for the cognitive projection differs between HC and AD participants (*p*=0.0053), with a faster age-related decline across the HC participants. Of note, the cognitive projection is significantly higher in the presence of both amyloid (*p*=0.0012) and the *APOE* ɛ4 allele (*p*=0.0039, Fig. 3c). The SW projection is not influenced by either of these factors.

### 3. Brain-behaviour modes in older adult ADNI participants

Application of PLS to the 807 healthy older adult participants from the ADNI cohort also yields a single robust mode (*p*<0.001, *z*=12.67). The cognitive projection loadings show effects comparable to the older AIBL cohort, covering all domains, with a markedly higher affinity for the memory domain (Fig. 4a). The brain anatomical projections are strongest over the posterior inferior temporal sulcus, the occipital lobe, the collateral fissure, the insula and the anterior inferior temporal sulcus (Fig. 4b; Supp. Fig. 1). Similar to the PISA data, using the CT as the brain measure yields a single mode that explains less *z*-covariance (*p*<0.001, *z*=4.88, Supp. Fig. 3).

**Figure 4:**
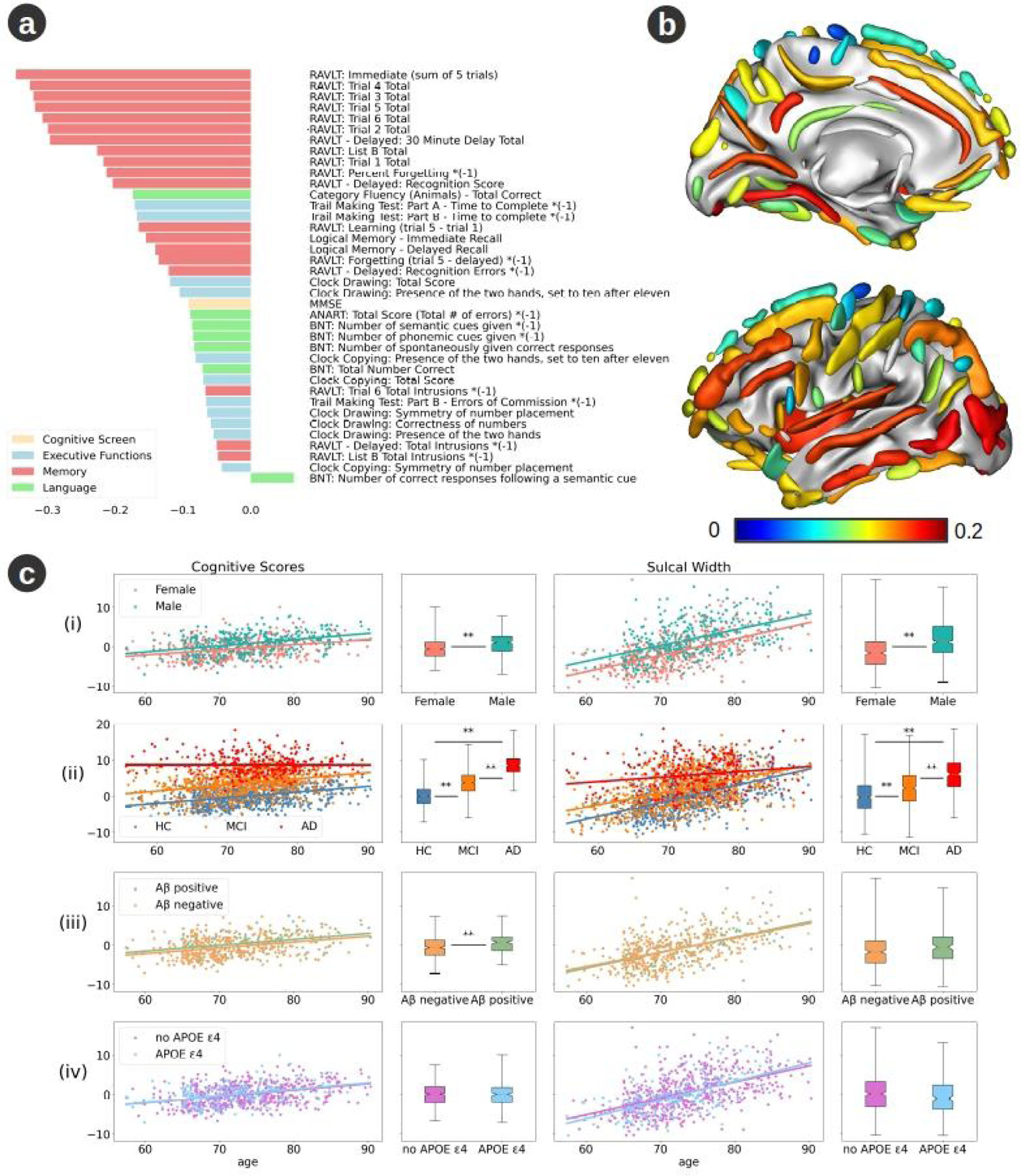
Robust PLS mode trained on ADNI healthy cohort. (a) Mean loading of all reliable cognitive scores. (b) Mean loading of all reliable cortical sulci. (c) Effect of age on ADNI brain-behaviour mode. Panels and legend as per Figure 2.

As with the PISA and AIBL data, both the cognitive and anatomical projections are correlated with the age of the ADNI participants (SW *p*=4.2e-68; cognition, *p*=3.0e-26) and differ for sex, when controlling for age (SW, *p*=7.9e-19; cognition, *p*=3.7e-10; Fig. 4c). When controlling for age and sex, both projections are significantly higher for AD than for MCI and for MCI compared to HC participants (Fig. 4c). Furthermore, the age-dependent slope differs between AD and HC (SW, *p*=1.8e-11; cognition, *p*=1.8e-8), between MCI and AD (SW, *p*=1.6e-6; cognition, *p*=1.6e-7) and for the brain projection between MCI and HC participants (SW, *p*=0.034; cognition, *p*=0.54). The cognitive projection is significantly influenced by the presence of amyloid (*p*=9.6e-4) but not by the *APOE* ɛ4 allele (*p*=0.31). The SW projection is not influenced by either of these factors (amyloid, *p*=0.64; *APOE*, *p*=0.56) (Fig. 4c).

### 4. Brain-behaviour modes across healthy and clinical participants

We next included the PISA clinical participants (with MCI or AD) alongside the healthy PISA participants, effectively increasing the cognitive variance into the clinically impaired range. Using SW as the anatomical measure and regressing out age and sex, yields a single robust PLS mode (1st mode, *p*<0.001, z=16.3; 2nd mode, *p*>0.99). Likewise using CT as the anatomical measure yields a single mode but with less covariance explained (1st mode, *p*<0.001, z=10.6; 2nd mode, *p*>0.99; Supp. Fig. 4).

Compared to the PLS model trained only on the PISA HC data, the robust mode including clinical participants loads with greater affinity onto the memory tests (Fig. 5a). The brain projection loads most strongly with the SW of the posterior inferior temporal sulcus, the sub-parietal sulcus, the posterior lateral fissure, the superior temporal sulcus and the occipital lobe (Fig. 5b; Supp. Fig. 1). The cognitive projection is higher for healthy amyloid positive participants (*p*=0.0027) but is not significantly influenced by the presence of *APOE* ɛ4 allele (*p*=0.28). The brain projection is not significantly influenced by the presence of amyloid (*p*=0.99) or the presence of *APOE* ɛ4 allele (*p*=0.53).

**Figure 5:**
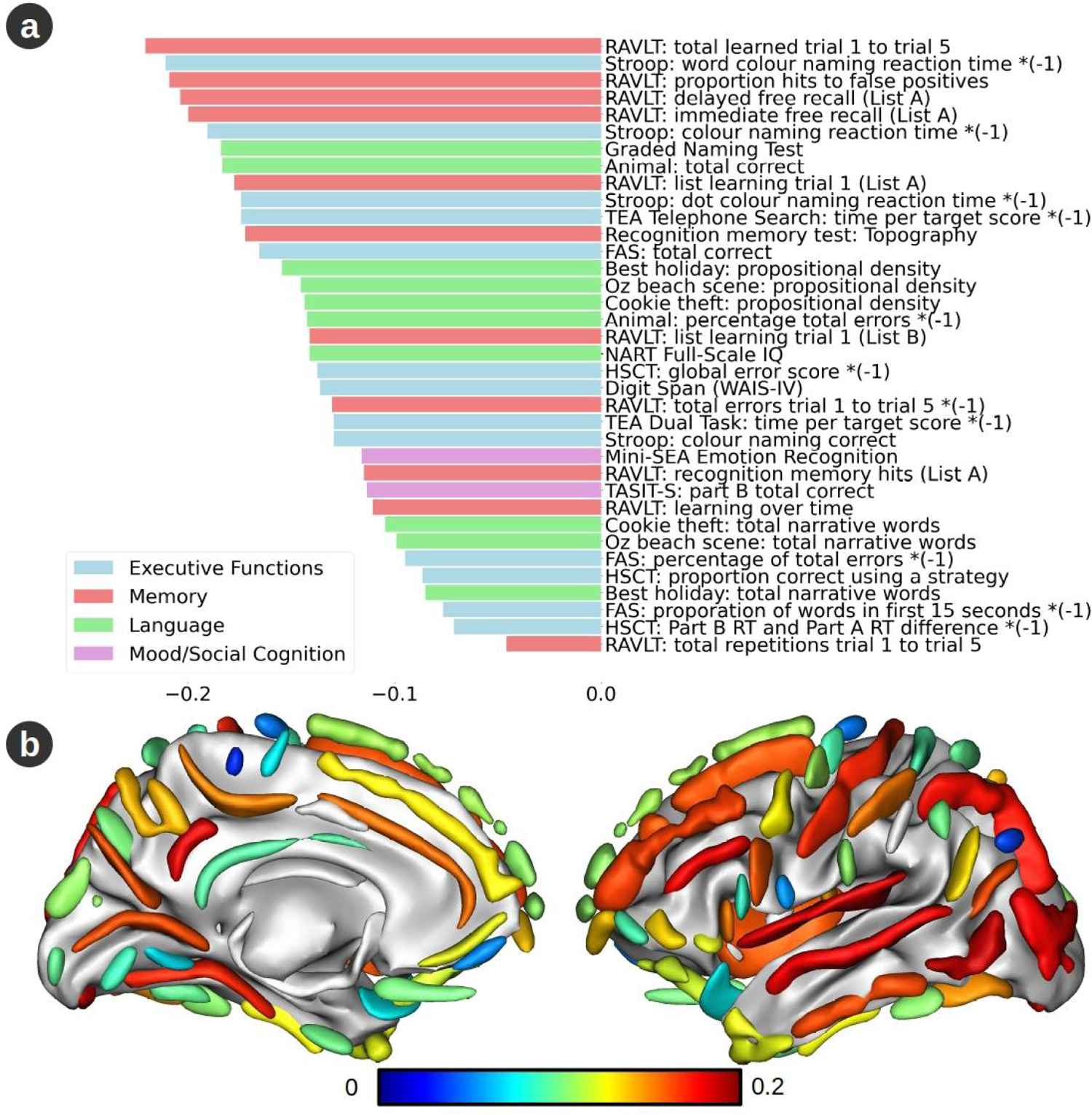
Robust PLS mode trained on the PISA cohort with both health (HC) and clinical (CC) participants. (a) Mean loading of all reliable cognitive scores. (b) Mean loading of all reliable cortical sulci.

Although the exact composition of the cognitive tests across the three studies differs, it is notable that the inclusion of clinical participants in the PISA PLS model biases the cognitive loading toward memory function, de-emphasizing executive function. Comparing the anatomical loadings across the cohorts (Fig. 6) shows that they bear stronger similarity when comparing the PISA model, including clinical participants, with the healthy AIBL/ADNI models, than when comparing the PISA model, including only (younger) HC participants, with the same AIBL/ADNI models. This suggests that healthy ageing centred in the eighth decade mirrors neurodegenerative changes occurring in the seventh decade rather than representing an extension of healthy ageing at that time.

**Figure 6:**
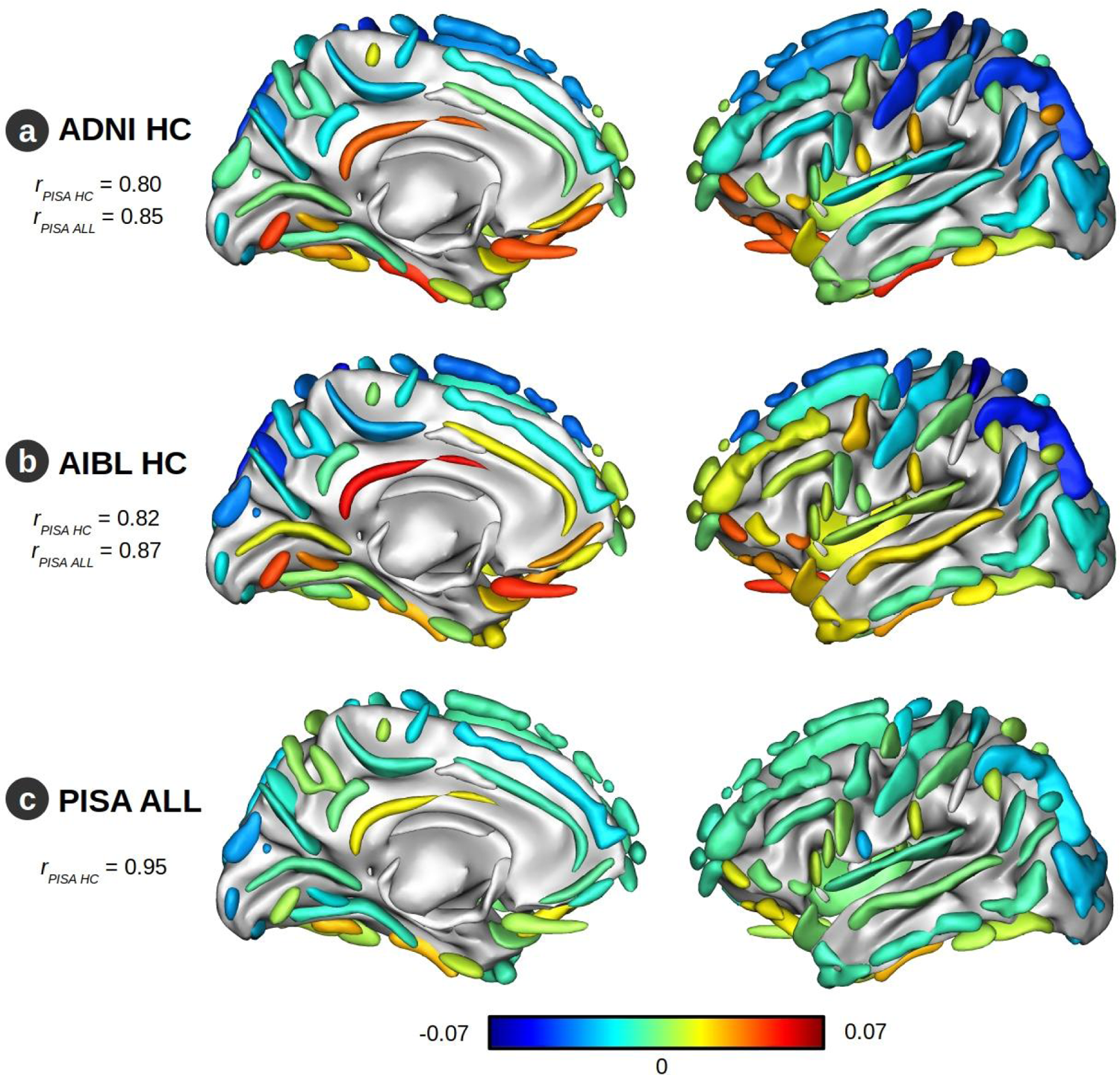
Brain loadings correlations and differences. (a) Differences in the SW loadings between the healthy ADNI cohort (ADNI HC) and the healthy PISA cohort (PISA HC). Positive values imply that the SW loadings of PISA HC are lower than those of ADNI HC. (b) Corresponding differences between the AIBL HC and PISA HC. (c) Difference between PLS trained on all PISA participants, after regression out of age and sex (PISA ALL) and healthy participants (PISA HC). The Pearson correlation coefficients (*r*) are shown between the brain loadings of the specified cohort (ADNI HC; AIBL HC; PISA ALL) and those of PISA HC (*r_PISA HC_*) and PISA ALL (*r_PISA ALL_*).

Comparing the anatomical loadings directly shows that healthy aging in the eighth decade (AIBL, ADNI) affects the fronto-temporal regions more and the occipito-parietal regions less than healthy aging (Fig. 6a,b) or neurodegenerative changes in the seventh decade (Supp. Fig. 5).

### 5. Influence of hippocampal volume on cognition

Considerable prior work has focussed on the role of changes in hippocampal volume in ageing and neurodegeneration (Fotuhi et al., 2012; Killiany et al., 2002; Pennanen et al., 2004). We hence sought to benchmark the relative influence of hippocampal volume (HV) against cortical sulcal width on cognition in the PISA cohort. Complimenting the list of cortical SWs by including left and right HV has negligible impact on the ensuing robust mode. Specifically, this mode is similar to the mode derived without inclusion of the HVs, with no significant impact on the cognitive projection (2.5 & 97.5 bootstrap percentiles: left HV {-0.03, 0.04}; right HV {-0.02, 0.06}).

Application of the PLS only on the HVs yields a single robust mode that is almost exclusively correlated with the memory tests (Fig. 7a) but explains relatively little cognitive covariance (*p*=0.050, *z*=1.73). In contrast to SW, the HV projections are not correlated with age (HV, *p*=0.087; cognition, *p*=0.0023), do not differ by sex (HV, *p*=0.26; cognition, *p*=8.1e-5) and do not differentiate MCI from AD participants (HV, *p*=0.76; cognition, *p*=0.038). However, the mode is significantly influenced by amyloid status (HV, *p*=0.037; cognition, *p*=5.5e-5; Fig. 7b) and does differentiate HC’s from MCI. Thus, amyloid accumulation in healthy subjects seems to be disproportionally related to HV loss rather than SW increase. However, we found no significant difference in the age-dependent effect on HV projections between Aβ+ and Aβ-participants (HV, *p*=0.75; cognition, *p*=0.18). Although both SW and HV projections show differentiation of HC from AD and MCI, SW projection performs a better differentiation than HV and their combination performs even better than either considered alone for differentiating HC, MCI and AD (Fig. 7c,d; Supp. Fig. 6).

**Figure 7:**
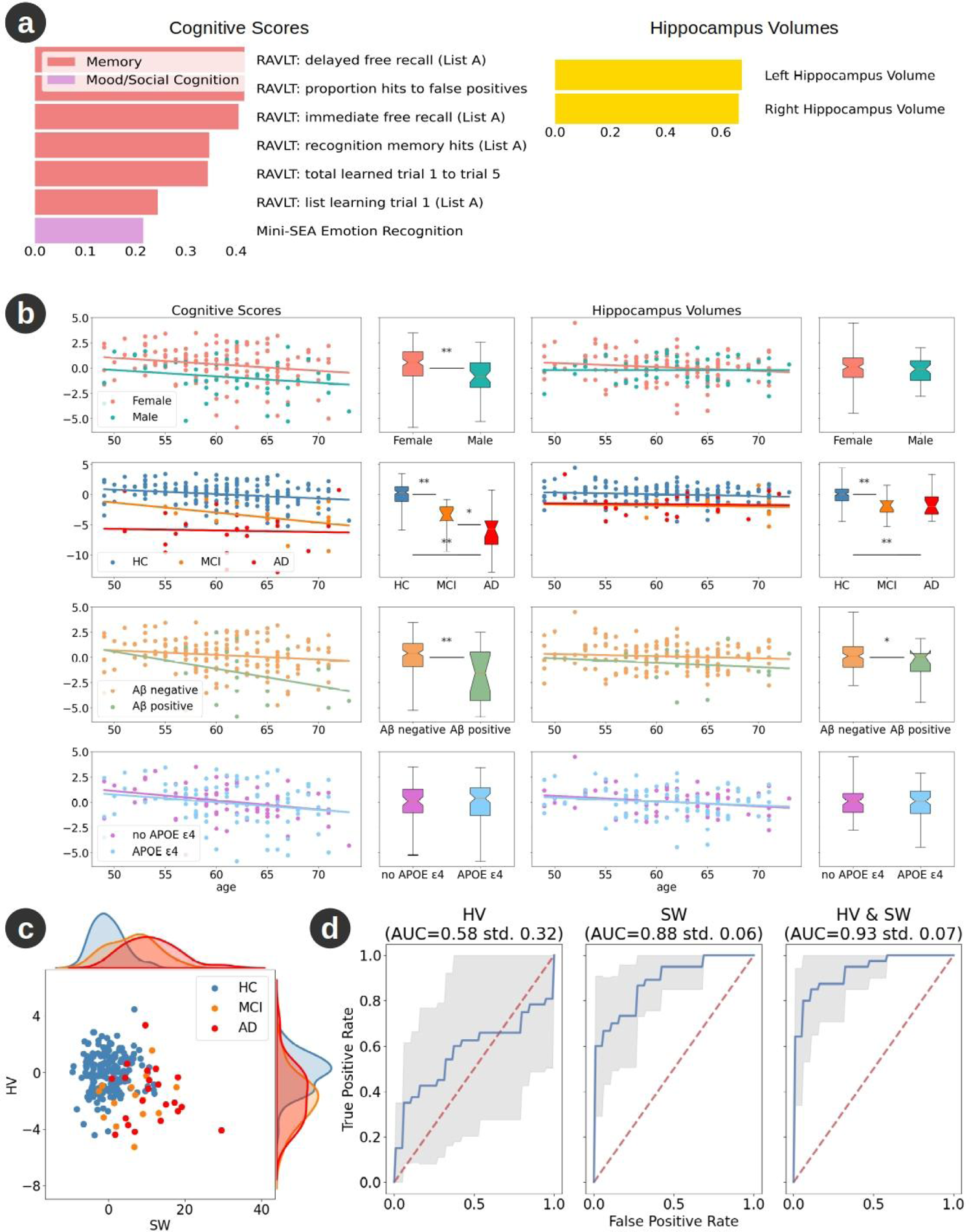
PLS mode trained on the healthy PISA cohort using hippocampal volume (HV) as the anatomical measure. (a) Mean loading of all reliable cognitive scores and of left and right HV. (b) Effect of age on HV-derived PISA brain-behaviour mode. Panels and legend as per Figure 2. Note that greater expression of this mode (more positive values) coincides with better performance of the corresponding cognitive loading (i.e. better performance on memory tests) and larger hippocampal volumes. (c) Comparison of hippocampus volume (HV) and sulcal width (SW) projections between PISA healthy cohort (blue) and participants with MCI (orange) or AD (red). Larger sulcal width covaries with smaller hippocampal volumes across participants. (d) Area under the ROC curve (AUC) after training a linear support vector machine to classify healthy and clinical participants, using a stratified 10-fold cross validation, based on the HV and/or the SW projections. The SW mode more accurately differentiates between HC and CCI participants than the HV-derived mode. The use of both SW and HV projections provides a slightly better differentiation between the two cohorts. Note that these classification models are based on anatomical measures only. Supplementary Figure 6 shows the results when age, sex, education and APOE status are included.

### 6. Healthy ageing and neurodegeneration

Although the preceding PLS modes from all three cohorts do not explicitly model age affects, they do all covary strongly with age – reflecting the strong influence of ageing on cognitive and anatomical variance across the age-range of these cohorts. These modes comprise multi-domain poorer cognitive performance and widespread wider sulci that also reflect individual variability. This raises the question of whether the specific composition of these modes are implicitly optimized to covary with age, or whether any linear weighting of poor cognition and wider sulci would perform comparably well. To test this, we randomly permuted the (SW and cognitive) mode features, producing surrogate PLS modes comprised of linear combinations of randomly chosen features. Performing this permutation 1000 times yields a reference distribution for non-specific anatomical and cognitive variability across our cohorts.

These tests show that the PLS trained on HC’s in each cohort always return the optimal combination of cognitive and SW weights that covary with age in the HC cohort, when benchmarked against randomly chosen features (Fig. 8a,b). Given that PLS is unsupervised, this finding suggests that age effects across the cohorts are stronger than inter-individual effects, and that (given the loadings differ) that these age effects differ between midlife and older adulthood. The age effects - trained on the HC participants within each cohort - also generally predict age effects in the MCI and AD participants, noting that in some of the cohorts, the age-effects in the clinical groups were reduced (such as AD in ADNI).

**Figure 8:**
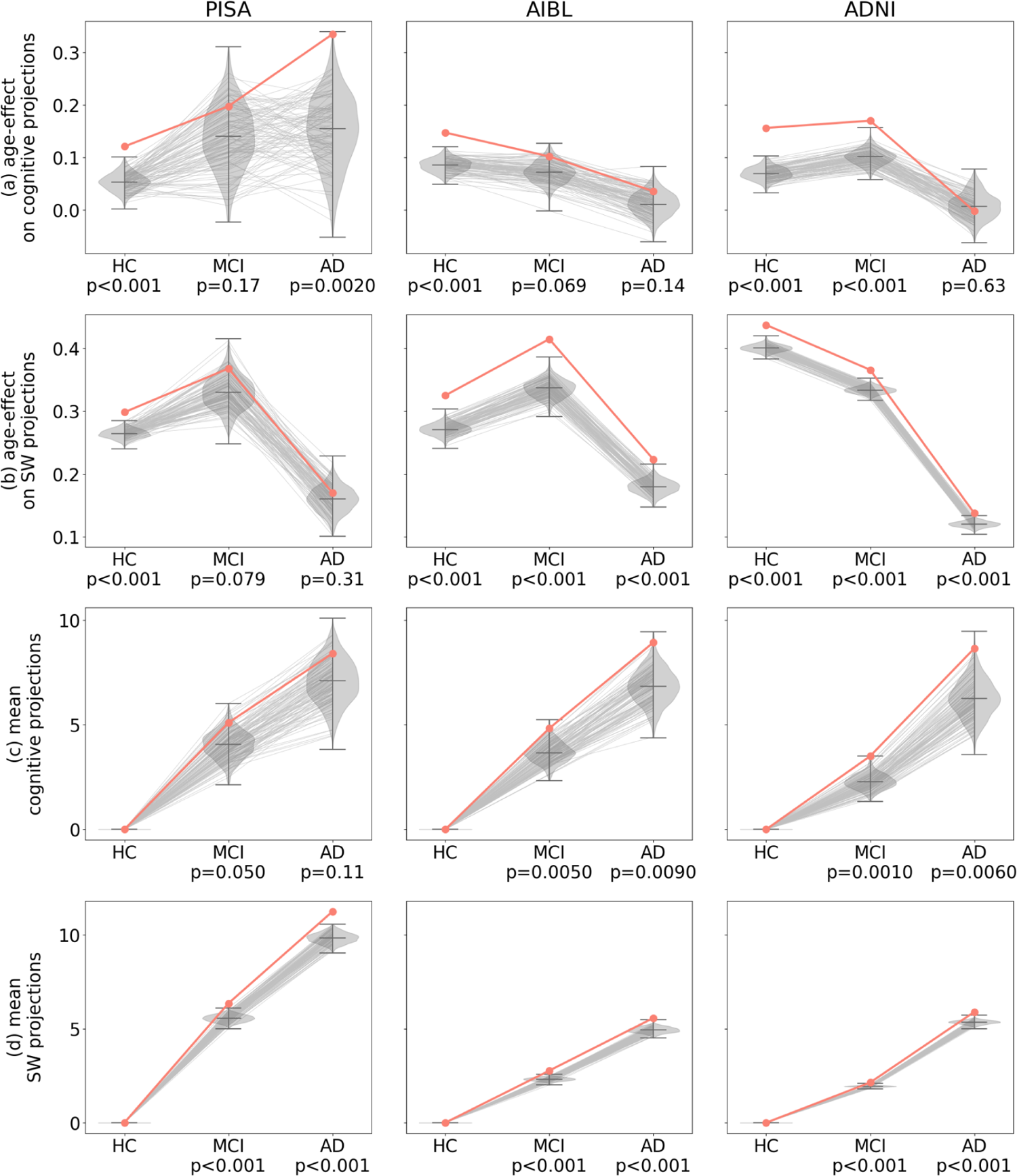
Specific versus non-specific age and diagnostic effects. Each column represents a different cohort. Results for the slope of the linear regression of the cognitive (a) and brain (b) projections against age (red) compared to randomly chosen features (grey). Results for the cognitive (c) and brain (d) projections for HC, MCI and AD participants, also benchmarked against randomly chosen features (grey). The violin plots represent the distribution of scores for 1000 permutation tests (bars represent the minimum, median and maximum value). The red line represents the original scores using PLS. The grey lines represent the scores for 100 representative permutations. The p-value is shown under the cohort label. Note that the p-value is not shown for the mean value of the healthy cohorts because the mean is always zero.

Finally, we tested whether the anatomical and cognitive features identified by performing PLS on the HC are optimized to identify out-of-sample MCI and AD, using the same permutation method (Fig. 8c,d). For all 3 cohorts, the exact combination of sulci always performs superior to randomly chosen sulci (Fig. 8d). For AIBL and ADNI, the unsupervised weights of cognitive scores are always superior to randomly selected cognitive tests. For the PISA cohort, the PLS mode trained on the HC ranks cognitive tests that usually, but not always, outperform randomly chosen tests (Fig. 8c).

## Discussion

In sum, this study demonstrates a single mode of covariation between cortical anatomy and cognitive function at midlife, and a single but substantially distinct mode in older adulthood. Age-related effects exert a strong influence on the expression of these modes, which also differ by gender, amyloid status and the presence of MCI or AD. As a result, brain-behaviour relationships in healthy data accurately and reliably predict MCI and AD diagnoses in matched clinical participants.

The presence of amyloid significantly impacts the cognitive projection, but not the SW projections, in all healthy cohorts. Given the substantial sample size (and hence power) of our study, the impact of amyloid on cognition but not SW suggests an impact on incipient neuronal integrity of cognitive relevance, but which does not (yet) impact large-scale cortical anatomy, even in the older cohorts. In contrast, for the hippocampal-derived PLS, amyloid influences both the anatomical (HV) projection and the associated (memory dominated) cognitive projection. This selective impact suggests that amyloid deposition leads to neuronal changes that deviate from those of healthy ageing (which influence many cortical regions and executive functions) leading to anatomically specific changes in the hippocampus and related cortical regions, such as the precuneus, with a disproportionate impact on memory (Farrell et al., 2018). However, somewhat paradoxically, we nonetheless observed that the pattern of cortical and cognitive variation across the healthy PISA cohort predicted out-of-sample AD and MCI substantially more accurately than expected by chance. This suggests a nuanced picture of targeted amyloid-related neurodegeneration, on the background of accelerated age-typical changes.

The presence of the ɛ4 allele of the *APOE* gene does not impact on the anatomical projections. It impacts cognitive projections for the older (AIBL) but not the mid-life (PISA) population. This may imply that its effect is less important, or even negligible, in mid-life adulthood. However, the PISA database has weaker power than AIBL, which may explain the lack of significant differences, particularly as the effect size in the AIBL data is small. Indeed, power analyses (alpha=0.05) for the results of the PLS trained on PISA HC show that the power is less than 20% when studying the influence of the *APOE* ɛ4 allele (SW, power=18%; cognition, power=19%).

Age-related changes depend on the age range studied. For the older populations studied (AIBL and ADNI), the influence of age is significantly more pronounced for healthy participants (independently of amyloid level) than for those with MCI or AD. This may be because participants with dementia present advanced brain ageing (Cole and Franke, 2017; Franke and Gaser, 2012), with a ceiling effect that mitigates the additional effect of chronological age. Conversely, for the mid-life population studied (PISA), age-related changes on cognition are weaker for healthy participants than for those with MCI or AD, and similarly for amyloid-negative compared to amyloid-positive healthy participants. Therefore, at mid-life, the early stages of dementia and amyloid accumulation accelerate cognitive ageing, whereas for older participants, dementia has already degraded cognition, overwhelming age effects in a cross-sectional study.

The composition and ranking of the cognitive domains varies across cohorts. Executive functions express the strongest brain-cognition covariance across the healthy mid-life (PISA) cohort, whereas performance on memory tests exerts the greatest influence when also incorporating individuals with early AD (PISA ALL) and in the older (ADNI, AIBL) cohorts. This suggests that mild to moderate brain neurodegeneration preferentially impacts memory while healthy mid-life brain ageing rather impacts executive functions. This questions the current trend to focus on memory tests to study ageing and AD. Although combining HV with SW does not impact the observed covariation mode, it is interesting to note that the HVs-cognition mode loads almost exclusively at mid-life with the memory tests, showing the importance of the brain measurement choice. For the older healthy populations studied (AIBL and ADNI), memory function exerts a dominant effect over the cognitive projections, similar to mild to moderate neurodegeneration at mid-life (PISA ALL).

Similar to cognition, the anatomical projections vary across cohorts. SW loads heavily on regions classically susceptible to ageing (prefrontal cortex, insula, superior parietal gyri, central sulci, cingulate sulci, calcarine cortex) (Burke and Barnes, 2006; Good et al., 2001; Salat et al., 2004) or by early progression of AD (temporal lobe, posterior cingulate, retrosplenial cortex) (Buckner, 2004; Killiany et al., 1993). Comparing AIBL/ADNI HC to PISA HC, suggests that fronto-temporal regions are more strongly involved with cognitive changes in the older population, while occipito-parietal regions are more involved in the mid-life population. Similarly, occipito-parietal regions appear more relevant for PISA HC than when also including the CC. The distinction between fronto-temporal and occipito-parietal regions in explaining cortical aging in relation to cognitive decline has recently been highlighted in (Cox et al., 2021), showing that fronto-temporal regions explain more of cortical ageing in the eight decade of life than occipito-parietal regions. Our study also supports this finding and suggests that the occipito-parietal regions explain relatively more of the healthy cortical ageing in the seventh decade of life.

Among the different brain measurements studied, sulcal width covaries more strongly with cognitive scores than cortical thickness or hippocampal volumes. Moreover, local anatomical contributions are specified less precisely when using CT than SW. As previously suggested (Bertoux et al., 2019), two advantages of SW may explain this: 1. Unlike CT, SW measurement does not depend on the grey-white matter boundary that blurs with age (Salat et al., 2009); 2. SW incorporates CT thinning as well as white matter reduction around the sulci. Compared to cortical SW, hippocampal volume at mid-life does not show an age effect and differentiates between the healthy cohort and the clinical cohort less accurately than the SW projection. Although surprising, note that the raw and ICV-corrected HV measures are also not significantly correlated with age and do not differentiate MCI subjects from AD subjects. The lack of a significant difference between MCI and AD might be due to the limited size of the clinical cohort. The non-correlation with age is intriguing and suggests that, unlike SW, HV is not impacted by direct associates of ageing (neural death) at mid-life but is vulnerable to pathological neurodegenerative processes, such as dementia and amyloid accumulation.

There are several caveats to our study. First, the healthy mid-life population studied (PISA HC) is enriched for people at high genetic risk of AD. Although we did not find a direct influence of the selection criteria (*APOE* status and AD-PRS) on the projections obtained, this selection policy could have influenced the results obtained. Second, each database used different cognitive tests, with a less focussed assessment of executive functions for ADNI. This could explain the slightly different results between AIBL and ADNI. Finally, this study focused on identifying patterns of covariation between cognition and structural MRI. Other MRI-derived measures, such as functional or diffusion MRI, could provide a complementary view (Damoiseaux, 2017).

## Methods

### 1. Participants

Cognitive, neuroimaging (MRI, PET) and genetic data from 1570 healthy adults (healthy cohorts, HC) and 1365 adults with MCI or AD (clinical cohorts, CC) were drawn from 3 multimodal databases; PISA, ADNI and AIBL (Fig. 1). All participants had a structural (T1-weighted) MRI scan and at least 50% of the cognitive scores available. The databases were sampled such that participants in each database were matched for age and sex (Supp. Table 1).

#### Prospective Imaging Study of Ageing (PISA)

The PISA cohort comprised a mid-life population enriched for high genetic risk of AD, derived from the Prospective Imaging Study of Ageing (PISA): Genes, Brain and Behaviour (Lupton et al., 2021). In addition to this genetically enriched sample, patients meeting formal criteria for MCI/AD across the same age range were recruited from local memory outpatient clinics (Lupton et al., 2021). We subsampled 190 healthy mid-age Australians (HC; mean age 61, range 49-73; 49 males) selected to be age- and sex-matched to 35 clinical participants with MCI or early onset AD (CC, mean age 63, range 51-72; 15 males). All data were acquired at a single site (Brisbane, QLD). The PISA study protocol, using written informed consent, has approval from the Human Research Ethics Committees (HREC) of the University of Newcastle, under the approval number H-2020-0439.

#### Australian Imaging, Biomarker & Lifestyle Flagship Study of Ageing (AIBL)

The AIBL cohort comprised older adults from the Australian Imaging, Biomarker & Lifestyle (AIBL) Flagship Study of Ageing (www.aibl.csiro.au). Data was collected by the AIBL study group. AIBL study methodology has been reported previously (Ellis et al., 2009). We selected 573 healthy participants ascertained at AIBL baseline (mean age 73, range 60-89; 255 males) and 191 age- and sex-matched participants meeting criteria for MCI or AD (mean age 74, range 59-85; 101 males). MRI data were acquired at two centres (Perth, WA; Melbourne, Vic) on 7 different Siemens scanners. Only scanners with greater than three participants were included.

#### Alzheimer’s Disease Neuroimaging Initiative (ADNI)

The ADNI cohort focuses on an older American healthy population, derived from the Alzheimer’s Disease Neuroimaging Initiative (ADNI) database (adni.loni.usc.edu). We selected 807 healthy participants (mean age 73, range 57-90; 356 males) and 1139 age- and sex-matched participants meeting criteria for MCI or AD (mean age 73, range 56-89; 554 males). Data was acquired at 67 different sites. Only sites with greater than three participants were included. The ADNI data have been curated and converted to Brain Imaging Data Structure (BIDS) format (Gorgolewski et al., 2016) using Clinica (Routier et al., 2021; Samper-González et al., 2018).

### 2. Structural and molecular neuroimaging

#### Brain

T1-weighted structural Magnetic Resonance Imaging (sMRI) data were used to study brain anatomy. Structural data were acquired using a 3D-MPRAGE sequence (see scanner information in Supp. Method).

Positron emission tomography (PET) data were used to quantify amyloid status (positive or negative) of all participants. For the PISA participants, PET data were acquired on a Biograph mMR hybrid scanner (Siemens Healthineers, Erlangen, Germany) with ^18^F-florbetaben, a diagnostic radiotracer which possesses a highly selective binding for β-amyloid in neural tissue (Fodero-Tavoletti et al., 2012; Rowe et al., 2008). Amyloid data for the ADNI and AIBL cohorts were assessed with one of the five following tracers: ^11^C-Pittsburgh Compound B, ^18^F-florbetaben, ^18^F-florbetapir, ^18^F-flutemetamol or ^18^F-NAV4694. The CapAIBL software (Bourgeat et al., 2018) was used to quantify each image into centiloids (CL) allowing the classification of the participants as amyloid positive (>20 CL) or negative (<20 CL). Non-negative matrix factorisation was used to improve centiloid robustness across tracers and scanners (Bourgeat et al., 2021). Amyloid data were not available on 17 of the HC PISA participants, 4 AIBL HC participants and 444 of the ADNI HC participants.

#### Cognition

Cognitive and mood assessments were conducted by trained neuropsychologists at all sites. Participants completed a battery of standardized tests selected to assess multidomain cognitive functions (memory, language, visuospatial, attention, processing speed, social cognition and executive function; Lupton et al., 2021; McKhann et al., 2011; Petersen et al., 2010). These tests were grouped into four categories (memory, language, executive functions and other, Supp. Table 2 & missing data in Supp. Fig. 7-9).

#### APOE ɛ4

*APOE* genotype was determined from blood-extracted DNA (Ellis et al., 2009; Lupton et al., 2021; Petersen et al., 2010). The *APOE* genotype was not available in 13% of the HC AIBL participants and 3% of the HC ADNI participants. Among the HC participants whose *APOE* genotype was available, the proportion of those with at least one ɛ4 allele of the *APOE* gene was 53% of the PISA participants, 29% of the AIBL participants and 32% of the ADNI cohort.

### 3. Data processing and modelling

#### Sulcal Width (SW), Cortical Thickness (CT)

The Morphologist pipeline of the BrainVISA toolbox (Borne et al., 2020) was used to extract local measures of brain anatomy (see Supp. Method for processing details). The pipeline was applied in a docker image as described in https://github.com/LeonieBorne/morpho-deepsulci-docker. This pipeline identifies 127 cortical sulci, 63 in the right hemisphere and 64 in the left hemisphere. We extracted both cortical thickness (CT) around each sulcus and the sulcal width (SW), which have both shown potential for the early detection of AD (Bertoux et al., 2019; Dauphinot et al., 2020). As in (Dauphinot et al., 2020), right and left hemisphere measurements are averaged when the same two sulci exist on each hemisphere, resulting in 64 unique measurements (see Supp. Fig. 10 for abbreviations and full labels). We then used ComBat, a technique adopted from the genomics literature (Johnson et al., 2007) and recently applied to cortical thickness data (Fortin et al., 2018), to combine and harmonize the sulcal measurements across acquisition sites while preserving age, sex and diagnosis covariates. Three sulci were missing in more than 50% of participants (S.GSM., F.C.L.r.sc.ant., S.intraCing) and were not used in further analyses.

For the PISA database, hippocampal volume was estimated separately for a comparative analysis. The volume of the right and left hippocampi were estimated using the CurAIBL platform (Bourgeat et al., 2015) and corrected by dividing by the intracranial volume (ICV) and multiplying by the average ICV of all participants (see Supp. Method for processing details).

#### Cognitive and mood scores

Each cognitive score was signed so that a more positive value indicated better performance (e.g. task accuracy) and more negative values indicated slower or worse performance (e.g. error rate, reaction time).

### Partial Least Square (PLS)

To study co-variation between cognitive and brain changes across mid- and older adulthood, we used partial least squares, a multivariate method that sorts modes of common variation according to their brain-cognition covariance explained. The Canonical Partial Least Square (PLS) approach (Wegelin, 2000), implemented in the Python library scikit-learn (Pedregosa et al., 2011), was used. Two datasets are given as inputs: the first contains sulcal anatomy measures (CT or SW of each sulcus), and the second comprises the individual cognitive tests. One latent variable is calculated for each dataset so that the covariance between them is maximized. The method iteratively calculates several pairs of latent variables. The first (principle) mode corresponds to the pair explaining the most covariance, and so on for the ensuing pairs. We refer to the pairs of latent variables as brain or cognitive projections. After assessing the robustness of each mode through permutation tests (see below), the contribution of each individual score (a specific cognitive test or sulci) to the shared variance is reflected in the corresponding loadings. Higher scores of these loadings correspond to better task performance and wider sulci, respectively.

As the default across the three data sets, we fitted the PLS models only to the healthy participants and permitted age effects to remain in the data. In auxiliary analyses on PISA participants, the effect of age and sex were first regressed out of each measure prior to fitting the PLS model on all participants (healthy, MCI and AD). To achieve this, a linear model with age and sex was fitted to predict each measure and the resulting prediction is then regressed, with PLS applied to the residuals.

For all analyses, missing values were replaced by the average of all participants (healthy or not) used to fit the PLS model. All measures were standardized by removing the mean of these same participants and scaling to unit variance before applying the PLS. The code for this study is available at https://github.com/LeonieBorne/brain-cognition-pisa.

### 4. Statistics

#### Permutation tests

PLS returns a series of modes, ranked by their covariance explained. Permutation tests were used to identify which of these modes were robust (Nichols and Holmes, 2002). These tests consist of randomly shuffling subjects from one of the data domains (in this case, the cognitive measures dataset) to perturb the specific association with the other domain (MRI). Then PLS is re-performed and the covariance is measured between each pair of latent variables. This test is repeated 1000 times. If the covariance of any given mode obtained from the empirical data is greater than 95% of those obtained from the first mode with permutation tests then the mode is considered robust. As in (Smith et al., 2015), we compared ensuing modes to the first mode because it extracts the highest explained variance in a null sample and can thus be viewed as the strictest measure of the null hypothesis (Wang et al., 2020). To assess the significance of the original data against the permuted distribution, we use the covariance z-scored by the null distribution.

#### Bootstrapping

We used bootstrapping to identify which individual measures have a significant impact on the PLS latent variables (Mooney et al., 1993). This approach consists of creating a new database of the same size by randomly selecting participants with replacement. PLS is then performed on the bootstrapped data and the loadings between each initial measure and the corresponding latent variable are calculated. This test is repeated 1000 times. If the 2.5 and 97.5 percentiles of the loadings obtained have the same sign, the measure (a specific sulcus or cognitive measure) is considered to have a statistically significant impact on the calculation of the latent variable.

### Statistical analyses

A series of statistical analyses were performed to assess the impact of specific risk factors for dementia on the latent variables. The impact of age was assessed using a two-sided hypothesis test, using the Wald Test with the t-distribution as the test statistic. To test whether any age-related effect differs between subgroups (diagnosis, sex, amyloid or *APOE* status), we used an analysis of covariance (ANCOVA) testing the interaction effect. The effect of sex (male, female) was evaluated using an ANCOVA, controlling for age. The effect of diagnoses (healthy, MCI, AD), amyloid status (positive, negative), and *APOE* status (presence of the ɛ4 allele or not) were evaluated using an ANCOVA controlling for age and sex. Because the PISA contains 16 twins, we removed one of each twin before using the ANCOVA to compare *APOE* status and controlled for age and sex. The PISA sample was enriched for high genetic risk of AD, including *APOE* ɛ4 positive as well as those in the highest quantile of risk for AD as defined by a polygenic risk score (PRS) combining common AD genetic risk variants with APOE ɛ4 omitted (Lupton et al., 2021). To control for any selection bias caused by APOE ɛ4 negative participants being enriched for other AD genetic risk variants, we also controlled the ANCOVA for the AD PRS used in the participant selection.

## Acknowledgements

PISA is funded by a National Health and Medical Research Council (NHMRC) Boosting Dementia Research Initiative - Team Grant [APP1095227]. Data collection and sharing for this project was funded by the Alzheimer’s Disease Neuroimaging Initiative (ADNI) (National Institutes of Health Grant U01 AG024904) and DOD ADNI (Department of Defense award number W81XWH-12-2-0012). ADNI is funded by the National Institute on Aging, the National Institute of Biomedical Imaging and Bioengineering, and through generous contributions from the following: AbbVie, Alzheimer’s Association; Alzheimer’s Drug Discovery Foundation; Araclon Biotech; BioClinica, Inc.; Biogen; Bristol-Myers Squibb Company; CereSpir, Inc.;Cogstate;Eisai Inc.; Elan Pharmaceuticals, Inc.; Eli Lilly and Company; EuroImmun; F. Hoffmann-La Roche Ltd and its affiliated company Genentech, Inc.; Fujirebio; GE Healthcare; IXICO Ltd.; Janssen Alzheimer Immunotherapy Research & Development, LLC.; Johnson & Johnson Pharmaceutical Research & Development LLC.; Lumosity; Lundbeck; Merck & Co., Inc.; Meso Scale Diagnostics, LLC.;NeuroRx Research; Neurotrack Technologies;Novartis Pharmaceuticals Corporation; Pfizer Inc.; Piramal Imaging; Servier; Takeda Pharmaceutical Company; and Transition Therapeutics.The Canadian Institutes of Health Research is providing funds to support ADNI clinical sites in Canada. Private sector contributions are facilitated by the Foundation for the National Institutes of Health (www.fnih.org). The grantee organization is the Northern California Institute for Research and Education, and the study is coordinated by the Alzheimer’s Therapeutic Research Institute at the University of Southern California. ADNI data are disseminated by the Laboratory for NeuroImaging at the University of Southern California.

## Author Contributions

L.B. analysed the data from PISA, AIBL and ADNI (extraction of sulcal measures from structural MRIs and application of PLS) and wrote the original draft of the paper, under the supervision of M.B.; M.K.L., C.G., G.A.R., A.C., P. Mosley, R.A., J.F., M.B. designed PISA and/or participated in data acquisition; P. Mosley and R.A. recruited participants for PISA; G.A.R. and A.C. acquired PISA cognitive data; A.F., P.B., J.F. analysed the PET scans to estimate amyloid status and PISA structural MRIs to measure hippocampal volumes; M.K.L. analysed PISA genetic data and provided *APOE* status and AD-PRS. All authors participated in reviewing and editing the paper.

